# Extracellular vesicle-localized miR-203 mediates neural crest-placode communication required for trigeminal ganglia formation

**DOI:** 10.1101/2023.03.14.532527

**Authors:** Yanel E Bernardi, Estefania Sanchez-Vasquez, Michael L. Piacentino, Hugo Urrutia, Izadora Rossi, Karina Lidianne Alcântara Saraiva, Antonio Pereira-Neves, Marcel Ivan Ramirez, Marianne E. Bronner, Natalia de Miguel, Pablo H. Strobl-Mazzulla

## Abstract

While interactions between neural crest and placode cells are critical for the proper formation of the trigeminal ganglion, the mechanisms underlying this process remain largely uncharacterized. Here, we show that the microRNA-(miR)203, whose epigenetic repression is required for neural crest migration, is reactivated in coalescing and condensing trigeminal ganglion cells. Overexpression of miR-203 induces ectopic coalescence of neural crest cells and increases ganglion size. Reciprocally, loss of miR-203 function in placode, but not neural crest, cells perturbs trigeminal ganglion condensation. Demonstrating intercellular communication, overexpression of miR-203 in the neural crest *in vitro* or *in vivo* represses a miR-responsive sensor in placode cells. Moreover, neural crest-secreted extracellular vesicles (EVs), visualized using pHluorin-CD63 vector, become incorporated into the cytoplasm of placode cells. Finally, RT-PCR analysis shows that small EVs isolated from condensing trigeminal ganglia are selectively loaded with miR-203. Together, our findings reveal a critical role in vivo for neural crest-placode communication mediated by sEVs and their selective microRNA cargo for proper trigeminal ganglion formation.

**SIGNIFICANCE STRATEMENT:** Cellular communication during early development plays a critical role. In this study, we demonstrate a unique role for a microRNA in cell-cell communication between the neural crest (NC) and placode cells (PC) during trigeminal ganglia (TG) formation. By utilizing loss and gain of function experiments in vivo, we demonstrate a requirement for miR-203 during cellular condensation to form the TG. We revealed that NC produces extracellular vesicles, selectively carrying miR-203, which is then taken up by the PC and regulates a sensor vector exclusively expressed in the placode. Taken together, our findings reveal a critical role in TG condensation for miR-203, produced by post-migratory NC and taken up by PC via extracellular vesicles.

## INTRODUCTION

Organogenesis requires the coordinated interaction of different cell types. In vertebrates, a good example is the interaction between neural crest (NC) cells and ectodermal placodes, two cell types of distinct embryonic origin that both contribute to the formation of cranial ganglia such as the trigeminal ganglion (TG). The TG is the largest ganglion in the head and is responsible for mediating sensation of pain, touch, and temperature in the face as well as innervating the sensory apparatus of the eye muscles and the upper and lower jaws (Breau and Schneider-Maunoury, 2015; Saint-Jeannet and Moody, 2014).

The NC, a population of multipotent cells specified in the dorsal neural tube of vertebrates embryos, migrates extensively and differentiates into diverse cell types including neurons and glia of the peripheral nervous system (Crane and Trainor, 2006). Placode cells arise from the surface ectoderm (Graham and Shimeld, 2013; Patthey et al., 2014), then ingress or invaginate into the cranial mesenchyme. Placode cells then interact with NC cells during condensation of the cranial ganglia, producing functional ganglia comprised of both neural crest- and placode-derived cells (Baker and Bronner-Fraser, 2001; Singh and Groves, 2016; Steventon et al., 2014). While both NC and placode cells contribute to trigeminal neurons, NC cells are the sole source of peripheral glia (Baker and Bronner-Fraser, 2001).

NC and placode cells are known to interact extensively; for example, *Xenopus* NC cells chase placodal cells by Sdf-mediated chemotaxis, and placodal cells are repulsed by a PCP and N-cadherin signaling mechanism (Theveneau and Mayor, 2013). Thus, precise cell-cell communication is required to facilitate mixing, proper positioning, aggregation, and differentiation of the forming cranial ganglia (Steventon et al., 2014). However, surprisingly little is known about the nature of interactions between NC and placode cells during ganglion formation.

In recent years, extracellular vesicles (EVs) have emerged as a novel mode of cell-to-cell communication. EVs are capable of transferring information from a donor cell to a recipient cell, leading to changes in gene expression and cell function (Abels et al., 2019; Crewe et al., 2018; Skog et al., 2008; Thomou et al., 2017; Valadi et al., 2007; Zomer et al., 2015). EVs are classified in three main types based on their size and biogenesis: exosomes (50–150 nm in diameter) that arise from multivesicular bodies (MVBs); ectosomes or shedding vesicles derived from the plasma membrane of cells through direct outward budding (100–1,000 nm in diameter); and apoptotic bodies that are released during apoptotic cell death (bigger than 1,000 nm in size, Théry et al., 2018). In particular, the subset of small EVs (sEVs) have been extensively studied in cancer cells, where exosomes and other EVs have been shown to drive multiple aspects of cancer metastasis, including the promotion of cancer cell motility, invasiveness, and premetastatic niche seeding (Sato and Weaver, 2018; Tao and Guo, 2020; Wortzel et al., 2019). sEVs contain proteins, RNAs, and the cargo of specific microRNAs (miRNAs) that can be transferred from a donor to a recipient cell, leading to changes in gene expression and cellular function in the receivers (Abels et al., 2019; Crewe et al., 2018; Skog et al., 2008; Thomou et al., 2017; Valadi et al., 2007; Zomer et al., 2015). miRNAs are a class of small non-coding RNAs that regulate gene expression at the post-transcriptional level mostly by binding to the 3’UTR of target transcripts and fine-tune their expression through degradation and/or translational repression (Bartel, 2009; Yates et al., 2013). They are known to play important roles in the fine regulation of gene expression during normal development, from gastrulation to complex organ formation. miRNAs specifically regulate epithelial plasticity by promoting both epithelial cell delamination and mesenchymal cell coalescence (Bernardi and Strobl-Mazzulla, 2021). In particular, we have previously shown that miR-203 is epigenetically repressed by DNA methylation in premigratory NC cells, thus allowing the upregulation of its target genes, *Snail2* and *Phf12,* which are essential for their epithelial-to-mesenchymal transition (EMT) (Sánchez-Vásquez et al., 2019; Strobl-Mazzulla and Bronner, 2012). Importantly, we observed that the miR-203 locus is rapidly demethylated during NC migration. At the end of migration, NC cells condense into different derivatives, like the cranial sensory ganglia. Thus, we hypothesized that ganglion condensation may require the reactivation of miR-203. Using the trigeminal ganglion as a model, we show that miR-203 is re-expressed in coalescing and condensed NC cells to regulate trigeminal ganglion assembly. Intriguingly, we find that miR-203 is required for trigeminal ganglion condensation. Further, we find that miR-203-containing sEVs are produced in NC cells, which are then taken up by placode cells in which the miRNA exerts its biological effect. Together, our findings reveal that miR-203 is upregulated in post-migratory neural crest cells and its transport into placode cells via small EVs is critical for trigeminal ganglion condensation.

## MATERIALS AND METHODS

### Embryos

Fertilized chicken eggs obtained from “Escuela de Educación Secundaria Agraria de Chascomús” were incubated at 38°C until the desired embryonic stage according to the criteria of Hamburger and Hamilton (HH) (Hamburger and Hamilton, 1992).

### DNA construct and electroporation

For *in ovo* electroporation, eggs were incubated horizontally until stage HH8 and vectors were injected on the neural tube and/or the trigeminal placode region by air pressure using a glass micropipette as described previously (McLennan and Kulesa, 2019). The miR-203 overexpressing, sponge and sensor vectors were described and characterized in our previous publication (Sánchez-Vásquez et al., 2019). pHluo_M153R-CD63-mScarlet (a gift from Alissa M. Weaver, Vanderbilt University School of Medicine, Nashville, TN, USA. (Sung et al., 2020) was amplified with two pairs of primers (pHluo-Fw: 5’-AAA ctc gag GCC ACC ATG GCG GTG GAA GGA G-3’; pHluo-Rev: 5’-AAA gct agc CTA GGA TCC CTT GTA CAG CTC GTC C-3’), digested (XhoI/NheI) and subcloned into the chick overexpressing pCIG vector. After injection, a platinum electrode is placed on each side of the embryo, and the chick embryos are electroporated with five pulses of 15 V for 50 ms on and 100 ms off intervals. After electroporation, eggs were then sealed with tape and incubated to the desired stages. Embryos were removed from eggs, placed in PBS and fixed in 4% PFA and utilized for immunohistochemistry or *in situ* hybridization protocol.

### In situ hybridization (ISH)

Embryos were fixed overnight in 4% PFA in PBS at 4°C and then utilized for whole-mount ISH for mRNA (Acloque et al., 2008) and miRNAs (Sánchez-Vásquez et al., 2019) following the previously published protocols. In both cases, the mRNAs and LNA probes were labelled with digoxigenin (Roche). Hybridized probes were detected using an alkaline phosphatase-conjugated anti-digoxigenin antibody (Roche, 1:2000) in the presence of NBT/BCIP substrate (Roche). After ISH, some embryos were re-fixed in 4% PFA in PBS, washed, embedded in gelatin, and cryostat sectioned at a thickness of 14-16 μm. Embryos were photographed as a whole-mount using a ZEISS SteREO Discovery V20 Stereomicroscope with an Axiocam 512 camera and Carl ZEISS ZEN2 (blue edition) software.

### Immunohistochemistry (IHC)

Whole embryos were fixed for 15 min in 4% PFA, washed in TBST (500 mM Tris-HCl, pH 7.4, 1.5 M NaCl, 10 mM CaCl_2_. and 0.5% Triton X-100) and blocked in 5% FBS in TBST for 1h at RT. Embryos were then incubated in mouse anti-Tuj1 (1:250; Covance) and/or mouse anti-HNK1 (1:10; supplied by the Developmental Studies Hybridoma Bank) overnight at 4°C diluted in TBST-FBS. Secondary antibodies were goat anti-mouse IgG2a Alexa Fluor 647 (1:500) and goat anti-mouse IgM Alexa Fluor 568 (1:500) for one hour at room temperature. After several washes in TBS-T, whole embryos or sections were mounted and imaged by using Carl ZEISS Axio observer 7 inverted microscope containing an Axiocam 503 camera and Carl ZEISS ZEN2.3 (blue edition) software.

### Extracellular vesicles isolation and characterization

Trigeminal ganglia were dissected and collected from ~80-100 HH17-19 stage embryos for each of the replicates required for the different characterization techniques. For TEM, a group of trigeminal ganglia were fixed in a 4% (v/v) glutaraldehyde solution in 0.1 M (v/v) cacodylate buffer, pH 7.2-7.4. Samples were gradually dehydrated with serial solutions of 50%, 70%, 80%, 90%, 95%, 100% acetone and then embedded in epoxyPolybed 8120 resin. Next, ultra-thin sections (~70 nm thick) were harvested on 300 mesh copper grids, stained with 5% uranyl acetate and 1% lead citrate, and observed with a FEI Tecnai G2 Spirit transmission electron microscope, operating at 120 kV. The images were randomly acquired with a CCD camera system (MegaView G2, Olympus, Germany).

For preparation of sEVs, trigeminal ganglia were treated with a solution of dispase and trypsin to obtain a single cell suspension. Samples were centrifuged at 10000 x g for 10 min and the supernatant was recovered to isolate sEVs. Then, the sample was filtered through 0.2 μm filter and then pelleted by centrifugation at 100,000 x g for 90 min to obtain an sEVs enriched fraction. As the protocol of EVs isolation include a filtration step using a 0.2 μm filter, the term sEVs (that include exosome and small size microvesicles) will be used throughout the text. sEVs were resuspended in PBS-DEPC containing a protease inhibitor cocktail (cOmplete™ ULTRA Tablets, Mini, EASYpack. Sigma). For characterization of particle quality size and abundance of the isolated sEVs nanoparticle tracking analysis methodology (NTA - Nanoparticle Tracking Analysis, Nanosight LM10 (Malvern™, U.K.)) was used, with 60-second readings performed in triplicate. In addition, a fraction of the sample was reserved for the isolation of small RNA.

### RNA preparation and RT-PCR

RNA was prepared from the trigeminal ganglion and sEV fraction using the RNAqueous-Micro isolation kit (Ambion) following the manufacturer’s instructions, and RNA was treated with amplification-grade DNaseI (Invitrogen). The reverse transcription reaction to obtain the cDNA was performed with the MystiCq® microRNA cDNA Synthesis Mix kit (Merck) and amplified by PCR using the following primers (miR-34-5p Fw: 5’-GCC GCT GGC AGT GTC TTA G-3’; miR-203 Fw: 5’-CCG GCG TGA AAT GTT TAG G-3’; and miR-UNI Rev: 5’-GAG GTA TTC GCA CCA GAG GA-3’).

### Explants culture

Neural crest and placode cells were electroporated separately *in ovo* at stage HH9. After treatment, the embryos were allowed to grow at 37°C and each tissues were dissected. A neural crest explant was placed next to a placodal explant in plates previously treated with fibronectin and containing DMEM medium supplemented with 10% fetal bovine serum and penicillin/streptomycin. The explant pairs were cultured at 37°C in 5% CO_2_ overnight. Cell culture experiments were imaged for two hours with a Zeiss LSM 980 inverted confocal microscope at 37°C in 5% CO_2_.

## RESULTS

### miR-203 is expressed in coalescing trigeminal ganglion cells

We first examined the expression pattern of miR-203 during the course of cranial NC migration, coalescence and condensation during trigeminal gangliogenesis (see scheme in figure 1A) by performing *in situ* hybridization (ISH) analysis at selected developmental stages in chick embryos. miR-203 was previously shown to be present in premigratory NC cells but down-regulated prior to their delamination from the neural tube (Sánchez-Vásquez et al., 2019). In agreement with this, we noted that mature miR-203, detected using locked nucleic acid-digoxigenin-labeled probes, was absent at HH13 from migrating HNK1 immunoreactive cranial NC cells (Fig. 1B). However, miR-203 expression was again noticeable at stage HH16 when cranial NC cells are coalescing at the site of trigeminal ganglion formation (Fig. 1C. Black arrowheads). Later at stage HH20 when the ganglion is almost fully condensed, signal was robust, particularly at the center of the lobe (Fig. 1D). Taken together, these data indicate that miR-203 expression is reactivated at the time of NC coalescence and condensation into ganglia, consistent with the intriguing possibility that miR-203 may be required for NC aggregation during trigeminal ganglion formation.

**Figure 1: miR-203 expression is activated during trigeminal ganglion formation. (A)** Schematic diagrams of a cross-section through the cranial neural tube (NT) illustrating the migration of NC and their coalescence and condensation with placode cells during trigeminal ganglion (TG) formation in chicken embryos. Transverse section of miR-203 after *in situ* hybridization using an LNA-DIG-labeled probe followed by HNK1 immunostaining to detect NC (red) cells at HH13 **(B)**, HH16 **(C),** and HH20 **(D)**. While miR-203 is absent from migrating NC cells, its expression reinitiates at the time of ganglion coalescence and remains present in the condensed ganglion. Black arrowheads denoted early coalescing cells expressing miR-203. NT, neural tube; TG, trigeminal ganglion.

### Overexpression of miR-203 generates ectopic aggregation of NC cells and a more condensed trigeminal ganglion

Given that miR-203 expression is down-regulated at the onset of NC migration and then re-expressed during trigeminal ganglion formation, we asked whether overexpression of miR-203 would accelerate the condensation process. To this end, we generated an overexpression vector containing pre-miR-203 (Sánchez-Vásquez *et al.,* 2019) which was electroporated into the right half of the neural tube at HH9. Embryos were then allowed to develop until HH15-16. The results show that excess miR-203 causes ectopic aggregation of NC cells, as identified by *in situ* hybridization against *Sox10,* compared with the control side or embryos treated with empty vector (Control OE) (Fig. 2A. Black arrowheads). In transverse section, ectopically aggregation of NC cells (Fig. 2A’. White arrowheads) and a more densely packed trigeminal ganglion (Fig. 2A’. Black arrowhead) are evident on the treated side compared to control side (Fig. 2B). These results suggest a possible role for miR-203 during trigeminal ganglion condensation.

**Figure 2:**
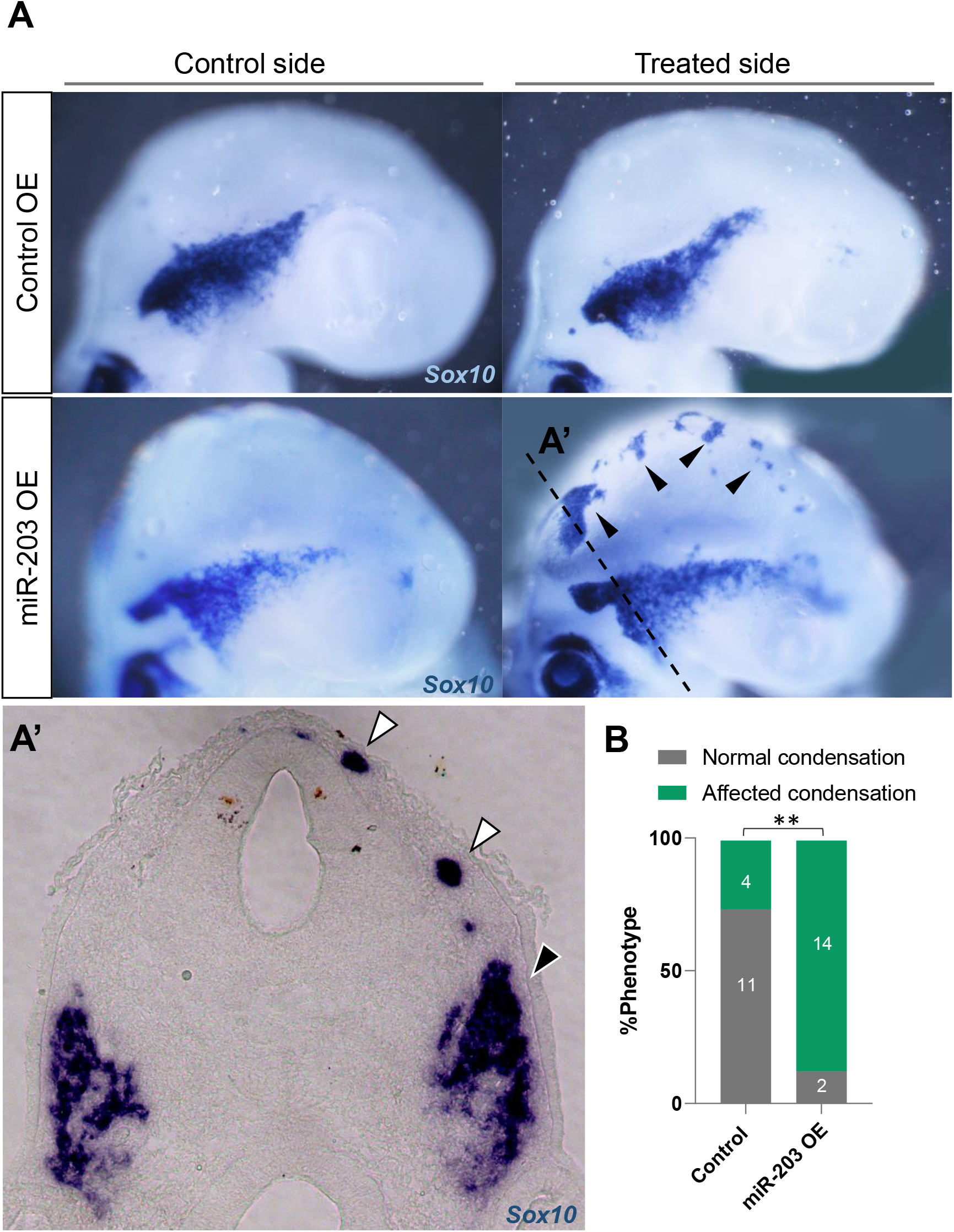
Overexpression of miR-203 promotes ectopic and premature NC condensation and enhanced aggregation into the trigeminal ganglion. **(A)** *In situ* hybridization for *Sox10* in HH15-16 embryos unilaterally electroporated with miR-203 (miR-203 OE) or empty control (Control OE) overexpression vectors. **(A’)** Cross section of miR-203 OE embryo at the level shown in A (dotted line) reveals an ectopic Sox10+ group of cells (white arrowhead) and a denser TG (black arrowhead) on the treated side. **(B)** Quantification of embryos showing a phenotype (normal versus affected condensation having ectopic condensation and/or denser TG) on Control and miR-203 OE. Numbers in the graph represent the numbers of analyzed embryos. \*\**P*=0.001 by contingency table followed by a χ^2^ test.

### Loss of miR-203 function in placode, but not NC, cells affects ganglion condensation

Given that miR-203 re-initiates during trigeminal condensation, we next asked whether its loss of function would disrupt proper ganglion formation. To test this possibility, we utilized a ‘sponge’ vector containing repeated miR-203 antisense sequences (miR-203 sponge) previously validated (Sánchez-Vásquez et al., 2019) to sequester endogenous miR-203. A sponge vector containing a miR-203 scrambled sequence (Scrambled sponge) was utilized as a control. The neural tube was electroporated at stages HH9 to target one side of the embryo, and embryos were then examined after the ganglia had condensed (HH17-18). Surprisingly, we failed to detect defects in the morphology of the trigeminal ganglion after the loss of miR-203 in NC cells, as visualized by ISH for *Sox10* (Fig. 3A).

**Figure 3:**
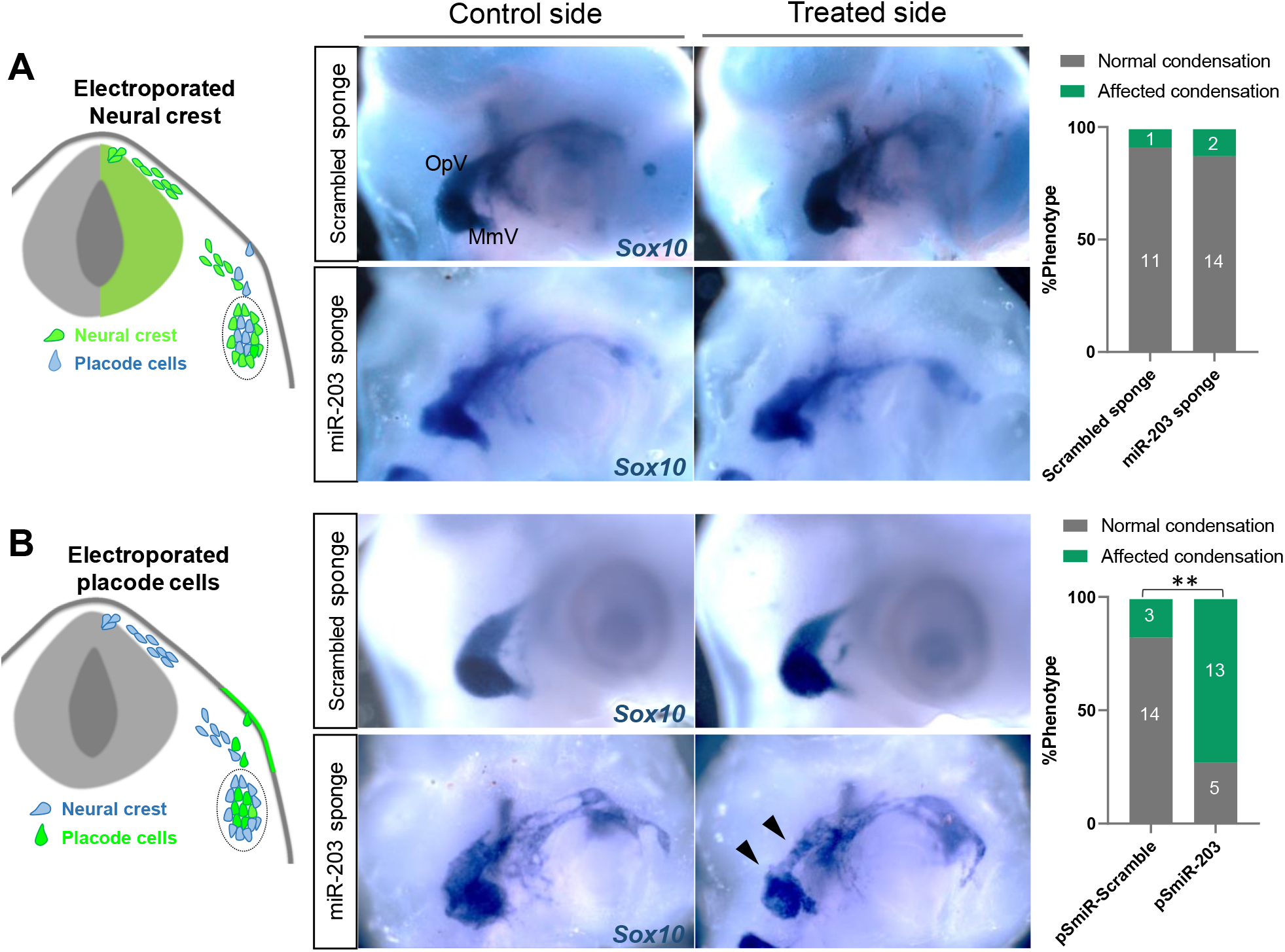
miR-203 loss of function in placode cells but not in the NC causes abnormal trigeminal ganglion condensation. **(A-B)** Electroporation scheme for region-specific knockdown of miR-203. Embryos at HH9 were unilaterally electroporated in the neural tube **(A)** or ectodermal placodes **(B)** with miR-203 or scrambled sponge plasmids and TG condensation was analyzed by ISH for *Sox10* at HH17-18. Bar graphs represent the quantification of embryos showing a phenotype (normal versus affected condensation having a less aggregated ganglion) on the miR-203 and Scrambled sponge treated embryos. Numbers in the graphs represent the numbers of analyzed embryos. \*\**P*=0.0015 by contingency table followed by a χ^2^ test.

The trigeminal ganglion has a dual origin from both NC and ectodermal placodal cells. Therefore, we next explored the possible functional role of miR-203 in the trigeminal placodes by electroporating the right placodal ectoderm at HH9 with the miR-203 or scrambled sponge plasmids. Intriguingly, the miR-203 sponge resulted in trigeminal ganglia that displayed a more loosely organized and less aggregated morphology than those in the non-injected side or observed in control embryos (Fig. 3B. Black arrowhead). The effect was statistically significant and more severe in the ophthalmic (OpV) than in the maxillo-mandibular (MmV) lobe; this is highly reminiscent of the phenotype observed after trigeminal ectoderm ablation (Shiau et al., 2008). Our findings raise the intriguing possibility that miR-203 is produced in the neural crest (donor cell) but exerts its biological effect in the placode cells (recipient cell). This is consistent with the putative role of placode cells as crucial mediators of NC condensation (Shiau et al., 2008), thus emphasizing the importance of cellular communication between the NC and placode cells to ensure correct aggregation in time and space to form the trigeminal ganglion.

### Extracellular vesicles produced by NC cells are taken up by placode cells

Small EVs, including exosomes, have been described as a novel mode of cell-to-cell communication (Valadi et al., 2007; Wortzel et al., 2019). In particular, sEVs can transport subsets of miRNAs, among other molecules, from a donor cell to a recipient cell, modifying gene expression and cell function in the latter (Chen et al., 2012; Hannafon and Ding, 2013; Villarroya-Beltri et al., 2013). With this in mind, we evaluated the production of sEVs during trigeminal ganglion condensation as a potential mediator of NC-placode communication. To this end, we first performed transmission electron microscopy (TEM) from dissected trigeminal ganglia at HH17. The results reveal cells in close contact within the condensing ganglion (Fig. 4A). These contain multivesicular bodies (MVB) with intraluminal vesicles located in the cytosol close to the plasma membrane (Figs. 4B-C). Some of MVB appear to be releasing exosomes-like structures into the extracellular space (Fig. 4B). Shedding vesicles protruding from the plasma membrane are also observed (Fig. 4C).

**Figure 4:**
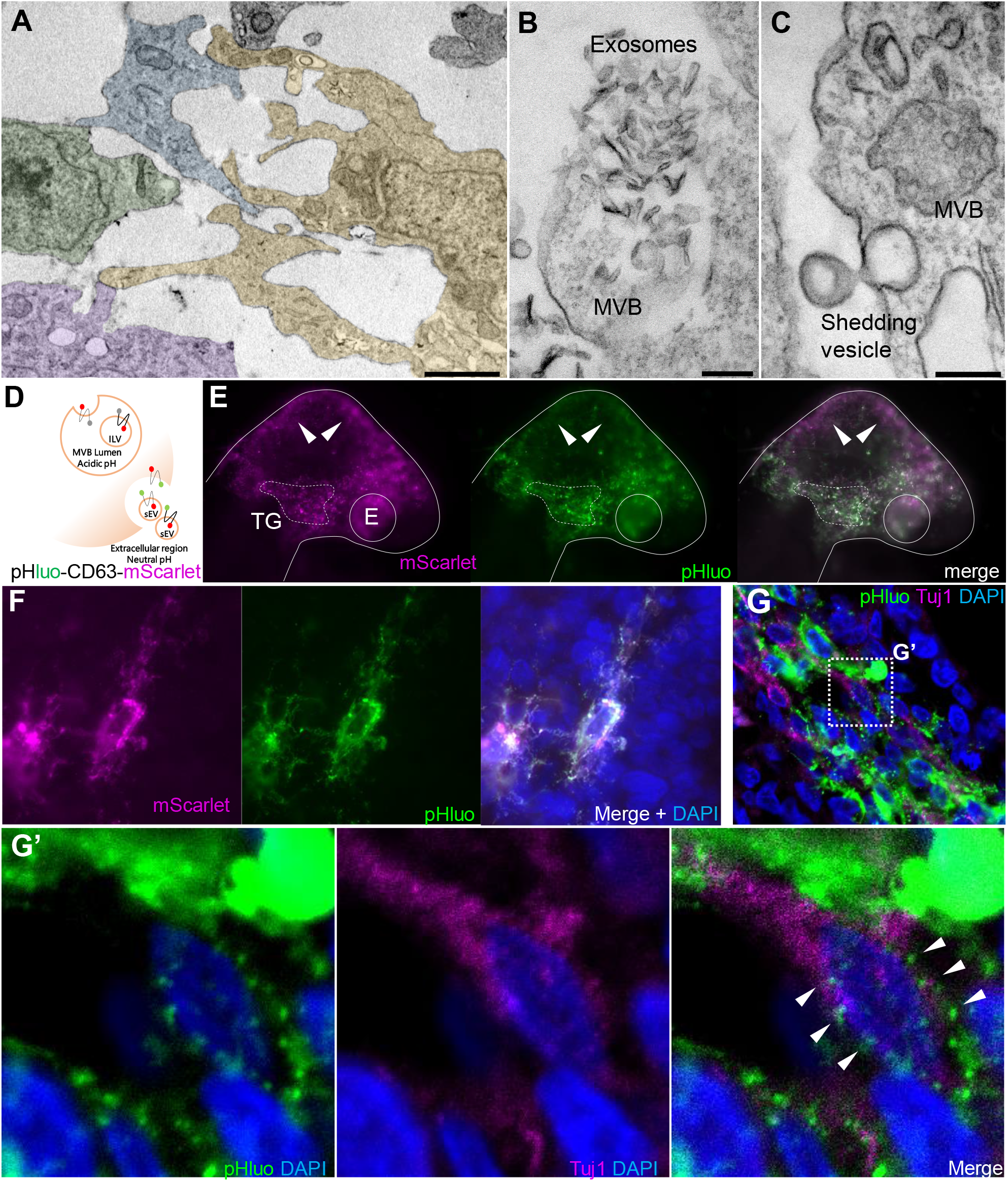
Extracellular vesicles are actively released by NC cells in the vicinity of placode cells during trigeminal ganglion condensation. Transmission electron microscopy from dissected trigeminal ganglia at HH16-17 shows that cells have many direct cell-to-cell contacts to each other **(A)**, the presence of multivesicular bodies (MVB) releasing exosomes-like structures into the extracellular space **(B),** and a shedding vesicle protruding from plasma membrane **(C)**. Bars: 2 μm (A), 200 nm (B and C). **(D)** Diagram of pHluo-CD63-mScarlet functional assay. **(E)** Embryos electroporated in the neural tube with pHluo-CD63-mScarlet at HH9- and visualized at stage HH16 exhibited GFP+/mScarlet+ fluorescence in the condensing trigeminal ganglion area (dotted line); in contrast, cells located in the midbrain are enrich in mScarlet+ (white arrowheads). TG, trigeminal ganglion; E, eye. **(F)** Migratory neural crest cells electroporated with pHluo-CD63-mScarlet exhibit intraluminal MVB (pHluo-/mScarlet+), extracellular/releasing EVs (pHluo+/mScarlet+) and deposit trails behind the cells. **(G)** Transverse section through the TG showing NC cells electroporated with pHluo-CD63-mScarlet and placode cells immunostained for Tuj1 (magenta). **(G’)** Zoom of box in G shows a placode cells (Tuj1+) with pHluo puncta incorporated into their cytoplasm (white arrowheads).

As a further confirmation that EVs produced by the NC cells are able to reach and be internalized by placode cells, we have adapted the pHluo_M153R-CD63-mScarlet vector (Sung et al., 2020) to work in chick embryos. This plasmid allows dynamic subcellular monitoring of exosome lifecycle, including MVB trafficking and exosome uptake. The plasmid contains a modified pH-sensitive GFP sequence (pHluo) inserted into the first extracellular loop of the tetraspanin CD63. Of note, pHLuo-CD63 does not fluoresce in the acidic endosomal pH of MBV; however, once exocytosed into the neutral pH of the extracellular environment, it emits a bright fluorescence. In addition, the plasmid is tagged with pH-insensitive red fluorescent protein (mScarlet) that allows the visualization of exosome trafficking (see scheme in figure 4D).

pHluo-CD63-mScarlet was introduced into the NC by electroporating into the neural tube of HH9^-^ chick embryos. After NC migration and TG condensation, we observed active vesicle release in the TG condensing region (dashed white line bordering the ganglia) compared with the midbrain region where only mScarlet fluorescence was detected (Fig. 4E, white arrowheads). Transverses sections through these embryos reveal mesenchymal migratory NC cells containing MVBs in acidic conditions (mScarlet only) and in neutral pH during the secretion process (mScarlet and pHluo positive) (Fig. 4F). Similar as previously described (Sung et al., 2020), we observed trails that may represent EVs deposition. Finally, to demonstrate the ability of NC cells to release EVs that can reach placode cells, sections of electroporated embryos with pHLuo-CD63-mScarlet were immunostained at HH16 for the neuronal marker Tuj1, which will mark placodally-derived neurons at this stage (Fig. 4G). Intermingled neural crest (pHluo+) and placode cells (Tuj1+) were detected in the region of trigeminal condensation, with the latter containing pHluo+ puncta incorporated into their cytoplasm (Fig. 4G’, white arrowheads). Taken together, these results demonstrate cellular interaction mediated by EVs between NC and placode cells *in vivo* at the time of trigeminal ganglion condensation.

To gain deeper resolution and reveal interactions in real-time, we next performed co-cultures with NC and placode explants from embryos electroporated with pHluo (pseudocolored yellow) and pCIG-mRFP (pseudocolored magenta), respectively (Fig. 5A). The explant pairs were cultured until migratory cells from both populations contacted one another at which time we generated time-lapse movies of the co-cultures. Live imaging revealed NC cells that were surrounded by numerous extracellular pHluo+ puncta (possibly EVs deposits), as well as trails (possibly migrasomes or retraction fibers) and cytonemes (Fig. 5B, Supplementary Movie 1). Interestingly, the cytoneme-like structures produced by migratory NC cells were in dynamic contact with placode cells (Fig. 5B’). In this sense, it has been shown that small EVs can travel along cytonemes and are released in close proximity to recipient cells during development (Chen et al., 2017; González-Méndez et al., 2017), raising the intriguing possibility that may be similar during neural crest-placode interactions.

**Figure 5:**
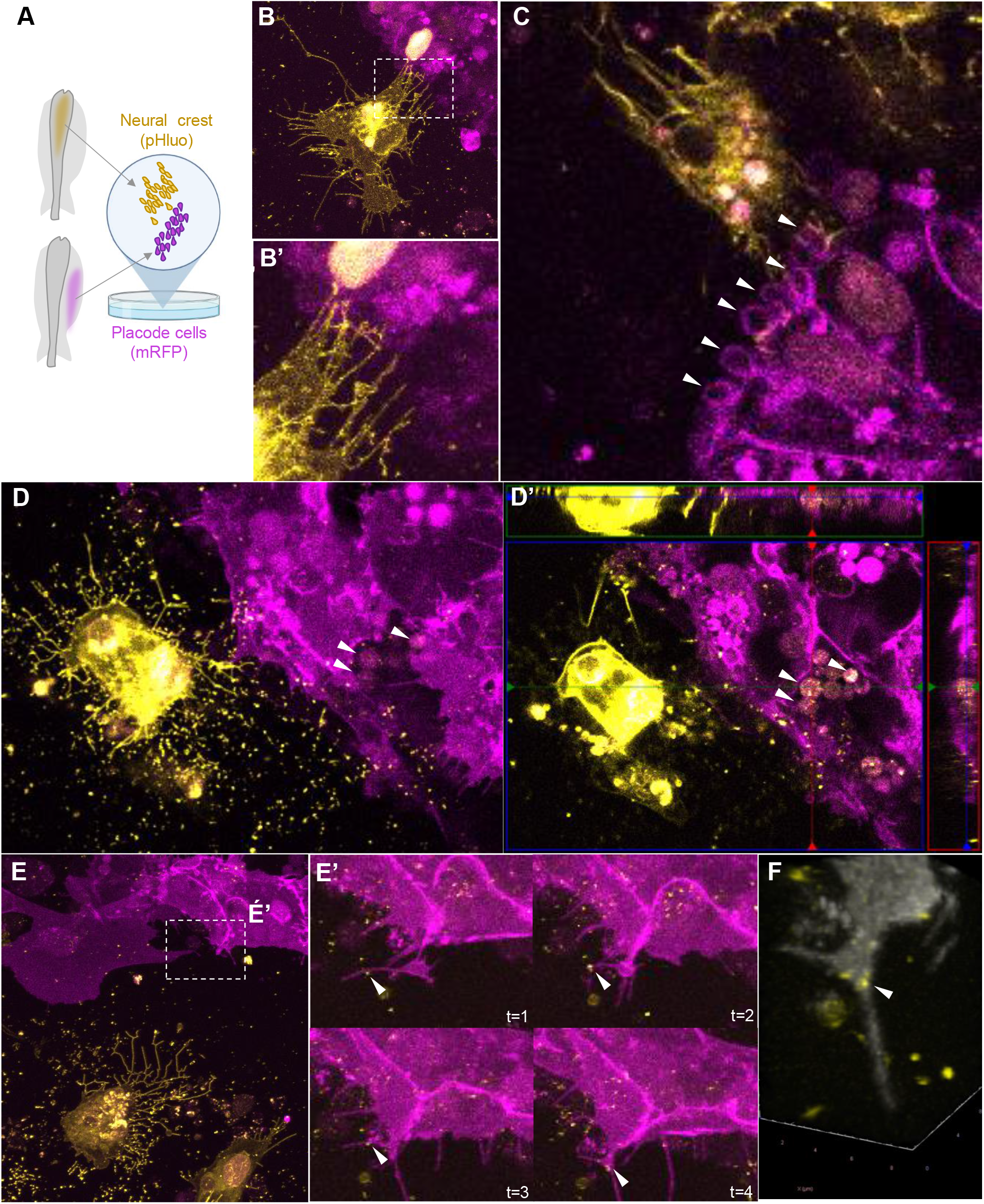
Extracellular vesicles produced by NC cells are engulfed by placode cells in explant co-cultures. **(A)** Scheme of a chicken embryo at HH9^-^ with neural tube electroporated with a pHluo (pseudocolored in yellow) and placode cells electroporated with membrane RFP (pseudocolored in magenta), respectively. The electroporated dorsal neural tube (mostly containing NC cells) and ectodermal trigeminal placode cells were dissected at HH9^+^ and HH10^-^, respectively, and co-cultured to examine their interactions. **(B)** NC cells (yellow) exhibiting trails (possibly migrasomes or retraction fibers) and cytonemes in very close contact with placode cells (magenta). **(B’)** Higher magnification of box in B showing the cytonemes from NC cells contacting placode cells. See also **Supplementary Movie 1**. **(C)** Placode cells showing vesicular structures like macropinosomes in their membrane (white arrowheads). See also z-stack in **Supplementary Movie 2**. **(D-D’)** NC cells expressing pHluo were surrounded by numerous extracellular pHluo+ puncta (possibly EVs deposits) and trails (possibly migrasomes or retraction fibers). Co-cultured placodal cells showing intra-cytoplasmic vesicular structures containing pHluo+ puncta (white arrowheads). See also **Supplementary Movie 3 and 4**. **(E)** Placode cell co-cultured with NC cells expressing pHluo produce filopodia that move toward the EVs deposits and engulf them. **(E’)** Zoom of box in E showing the time-lapse showing placode cells engulfing a pHluo+ puncta (white arrowhead). See also **Supplementary Movie 5**. **(F)** 3D reconstruction of placode protrusion observed in (E) demonstrating the internalized pHluo+ puncta (white arrowhead). See also **Supplementary Movie 6.**

It was previously shown that exosomes could be captured by filopodia or macropinocytosis events and endocytosed by recipient cells (Heusermann et al., 2016). We observed a similar event in which placode cells contained vesicular structures in their membrane (macropinosomes) (Fig. 5C, white arrowheads. Z-stack in Supplementary Movie 2) and cytoplasm (Fig. 5D-D’, white arrowhead. Time lapse and z-stacks in Supplementary Movie 3 and 4, respectively) containing pHluo puncta coming from co-cultured NC cells. Interestingly, we also noted that placode cells produced filopodia that move toward EVs deposits and engulf them (Fig. 5E-F, white arrowhead. Time lapse and 3D rotation in Supplementary Movie 5 and 6, respectively). These observations demonstrate that neural crest-produced EVs are internalized by placode cells.

### miR-203 produced in NC cells regulates translation in recipient placode cells

To determine whether miR-203 produced in NC cells can reach placode cells to exert a biological effect, we designed an experiment in which we drove overexpression of miR-203 and EGFP in the NC cells by electroporation into the premigratory NC. In the same embryos we electroporated the trigeminal placode in the ectoderm with a dual-colored sensor vector which expresses both nuclear d4EGFPn, containing two mature miR-203 recognition sites such that the miRNA can bind and affect protein translation (Sánchez-Vásquez et al., 2019), and mRFPn (see scheme in Fig. 6A). Embryos were electroporated at HH9- with the two vectors and allowed to grow until HH17. Transverse sections through the embryos were then immunostained for Tuj1 to identify the trigeminal ganglion cells (Fig. 6B). Although some of the placode cells reaching the condensing area were EGFP+/RFP+ (white arrowhead), some cells within close proximity to the NC (black arrowhead) were only RFP+ (dotted circles) (Fig. 6B’). This reduced EGFP expression is not visible in the non-migrating ectodermal cells where all the cells co-express both EGFP and RFP cells (white arrowheads in figure 6B). Importantly, embryos over-expression miR-203 displayed a significant reduction compared with controls when the EGFP/RFP fluorescence intensity ratios were quantified on placode cells (Fig. 6D).

**Figure 6:**
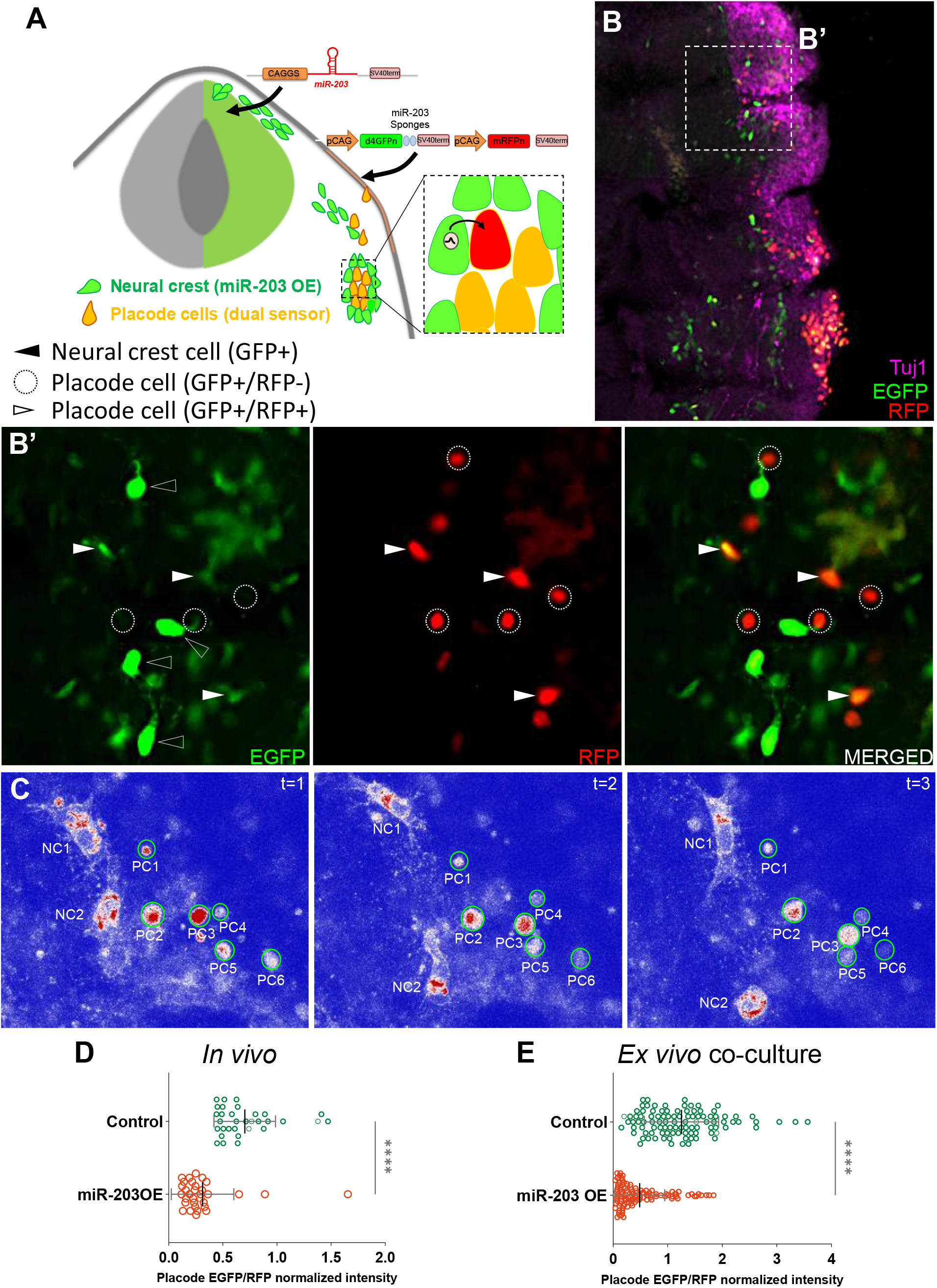
miR-203 generated in NC cells inhibits a sensor target electroporated in placode cells both *in vivo* and *ex vivo*. **(A)** Scheme of chick embryo with neural tube electroporated with miR-203 or empty control overexpressing vectors (with cytoplasmic EGFP) and placode cells electroporated with a dual-colored sensor vector (pSdmiR-203) containing two copies of complementary sequences to the mature miR-203. pCAG, Chick β-actin promoter; d4EGFPn nuclear-localized destabilized EGFP with a half-life of 4 h; mRFPn, nuclear-localized monomeric red fluorescent protein. Box of the condensing trigeminal ganglion indicates area of enlarged inset exemplifying the transfer of miR-203 from NC (green) to placode cells (orange because of the expression of both EGFP and RFP), thus affecting the EGFP translation (red cell). **(B)** Transverse section of HH17 embryos immunostained for Tuj1 (magenta), to identify the trigeminal ganglia with coalescing NC (electroporated with miR-203 OE vector) and placode (electroporated with the dual-colored sensor vector. See white arrowheads) cells. **(B’)** Zoom of box in B identifying migratory NC cells with cytoplasmic EGFP+ (black arrowhead) and placode cells with nuclear EGFP+/RFP+ (white arrowhead) or only RFP+ (dotted circles). **(C)** Time-lapse of co-cultured placode ectodermal cells (electroporated with the dual-colored sensor vector) and dorsal neural tube (electroporated with miR-203 OE) explants. Placodal cells (PC1-6) interact with NC cells (NC1-2) where the EGFP channel was pseudocolored (red>white>blue) to visualize the EGFP decay in placode cells over time. **See also Supplementary Movie 7**. **(D)** Scatter plot from *in vivo* experiments analyzing the EGFP/RFP intensity for individual placode cells reaching the condensing area at HH17 from miR-203 OE or Control electroporated embryos (cells counted in 2-5 sections per embryo, n=4 embryos from two independent electroporations). \*\*\*\**P*<0.0001 calculated using unpaired Student’s *t*-test. Error bars indicate standard deviation. **(E)** Scatter plot from *ex vivo* co-cultured experiment analyzing the EGFP/RFP normalized intensity for individual placode cells interacting with NC cells from miR-203 OE or Control electroporated embryos (100 cells, n=4 co-cultured explants for each treatment from 2 independent experiments). \*\*\*\**P*<0.0001 calculated using unpaired Student’s *t*-test. Error bars indicate standard deviation.

A similar experiment to that shown in figure 5A was performed where separate embryos were electroporated either with miR-203 or control over-expression vectors in the neural tube, and the dual-sensor vector was electroporated in the trigeminal placode. Explants from both dorsal neural tube and ectodermal placode were co-cultured *ex vivo* until migratory cells from both populations were in contact. LUT pseudocolored cells for EGFP intensity (RED>WHITE>BLUE) were analyzed by time-lapse demonstrating that placode cells (with nuclear EGFP+/RFP+) lose EGFP intensity over time when they are in close proximity to the NC cells (Fig. 6C. Supplementary Movie 7). In addition, EGFP/RFP intensity ratios in placode cells co-cultured with miR-203 over-expressing NC cells were quantified, and showed a significant decrease compared with controls (Fig. 6E).

Taken together, our *in vivo* and co-cultured explant results demonstrate that miR-203 produced in the NC reaches and suppresses translation in placode cells, thus supporting the idea that miRNAs may act as intercellular signals mediating proper neural crest-placode communication.

### Endogenous miR-203 is specifically loaded into sEVs produced during TG condensation

To determine the possibility that miR-203 may be loaded into EVs released in the context of trigeminal condensation, we performed two independent isolations of sEVs from ~80 trigeminal ganglia dissected at ~HH17. To demonstrate sEVs enrichment, we first analyzed the samples by nanoparticle tracking analysis (NTA) and transmission electron microscopy (TEM). The results show that an enrichment of small EVs with median size of ~80 nm in diameter (Fig. 7A) and the typical cup-shape of exosomes in the samples (Fig. 7B).

**Figure 7:**
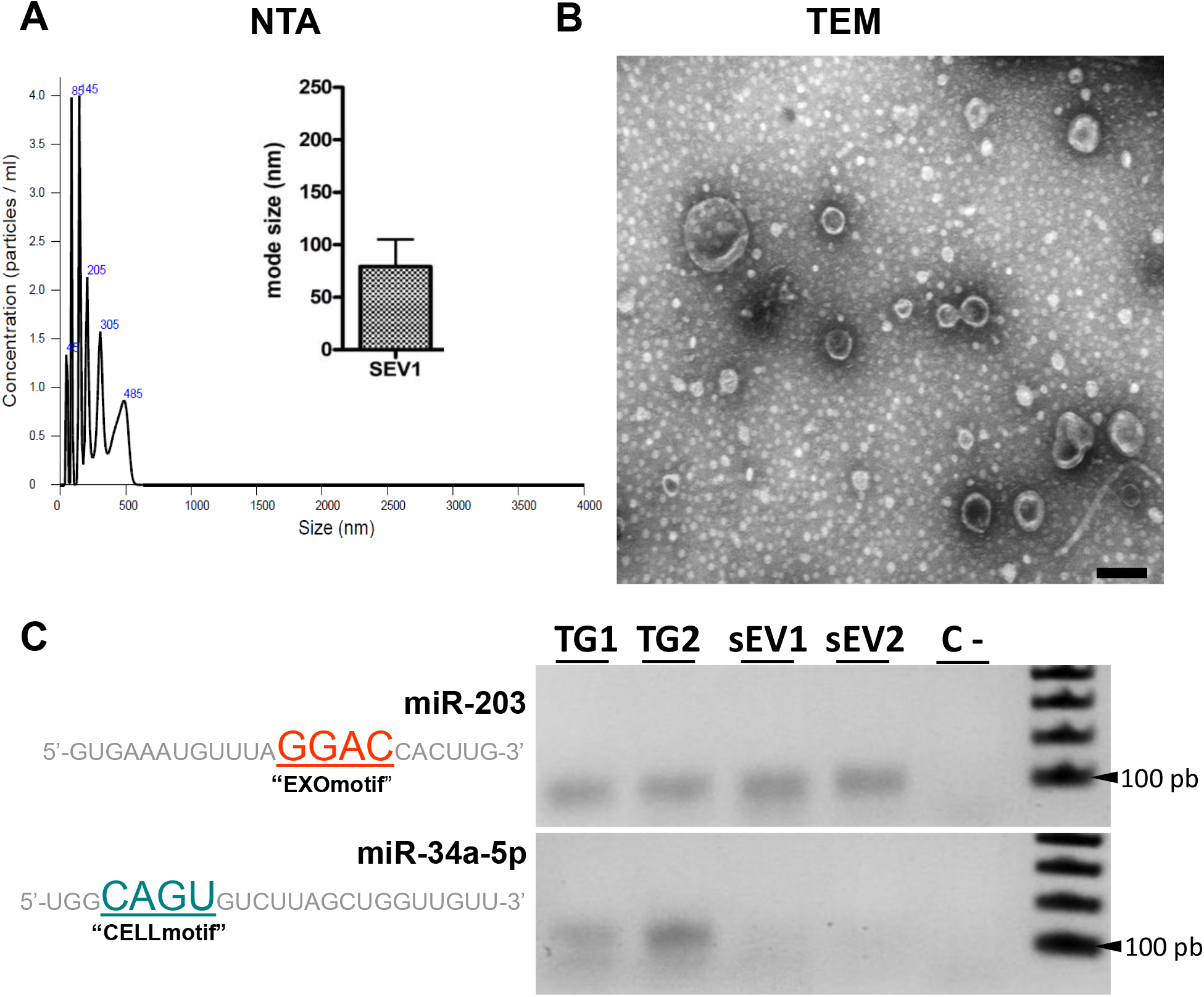
Endogenous miR-203 is selectively loaded into sEVs produced during trigeminal ganglion coalescence. **(A)** Graphs show particle concentration, size, and mode size using nanoparticle tracking (NTA) from enriched sEVs samples isolated from dissected trigeminal ganglia at HH17. **(B)** Transmission electron microscopy (TEM) image showing isolated exosomes after sEVs purified from dissected trigeminal ganglia. **(C)** RT-PCR from two independent whole dissected trigeminal ganglia (TG1 and TG2) and isolated sEVs from condensing trigeminal ganglia (sEV1 and sEV2) raised against the endogenous miR-203 (containing the EXOmotif) and miR-34a-5p (containing the CELLmotif).

Next, we isolated RNAs from these samples to determine if miR-203 is loaded into sEVs. It has recently been demonstrated that miRNAs are specifically loaded into sEVs or retained in the cells by RNA-binding proteins based on the presence of EXOmotifs or CELLmotifs, respectively (Garcia-Martin et al., 2022). To investigate whether this is the case during trigeminal ganglion condensation, we evaluated by RT-PCR the expression of miR-203 (having the EXOmotif GGAC) and miR-34a-5p (having the CELLmotif CAGU) in dissected TGs and isolated sEVs (Fig. 7C). The results show that miR-203 is detectable in both trigeminal ganglion samples (TG1 and TG2) as well as in the isolated extracellular vesicles (sEV1 and sEV2). In contrast to miR-203, and in agreement with the presence of the CELLmotif, miR-34a-5p was only detected in the TG and not in sEVs. These results demonstrate that endogenous miR-203 is selectively loaded into sEVs at the time of trigeminal ganglion condensation.

## Discussion

The development of the trigeminal ganglion involves a series of organized events to ensure its correct positioning, aggregation, and differentiation (Steventon et al., 2014), including intimate interactions between NC and placode cells. In our study, we demonstrate that this communication is at least partly mediated by the sEVs released by NC cells that are engulfed by and function within placode cells. Moreover, the selective cargo of miR-203 into sEVs, and possibly other miRNAs, acts as a signaling molecule required for neural crest-placode communication during their aggregation to form the trigeminal ganglion. While this interchange of sEVs between cell types has not previously been noted during development, it has been studied during tumor metastasis (Becker et al., 2016; Chang et al., 2021).

During formation of peripheral ganglia, mesenchymal migratory cells undergo several cellular and molecular changes. Our previous work demonstrated that the epigenetic repression of miR-203 is required for pre-migratory NT cells to initiate an epithelial-to-mesenchymal transition (EMT) (Sánchez-Vásquez et al., 2019). In this context, miR-203 may reflect a pro-epithelial or pro-aggregation cue. Consistent with this idea, reactivation of miRNAs that are characteristic of the aggregate state, such are miR-203, may be required for proper cell aggregation during organogenesis in early development, as well as during the formation of secondary tumors (Bernardi and Strobl-Mazzulla, 2021). The process of trigeminal ganglion aggregation has been likened to a mesenchymal-to-epithelial transition (MET) which is the reverse of EMT. However, placode and NC cells do not become completely epithelial during gangliogenesis, such that this transition may more appropriately reflect a mesenchymal-to-ganglionic transition (Lee et al., 2020). Together, our work has positioned miR-203 expression as a reversible mechanism mediating neural crest EMT during emigration and MET during trigeminal ganglion condensation.

Here we show that miR-203 functionality is not limited to the neural crest during EMT, but rather plays a later role in regulating condensation with placode cells, thus highlighting the importance of intercellular communication during trigeminal ganglion assembly. In this regard, it is well known that coordination between these cells is critical, and NC cells act as a guide and scaffold for the organization of placode cells to ensure proper assembly (Freter et al., 2013; Shiau et al., 2008) and the correct orientation of their projections to the central nervous system (Begbie and Graham, 2001; Freter et al., 2013).

Neural crest cells communicate with each other and with the extracellular matrix through cell contact, macropinocytosis, filopodia, and via gap junctions (Jourdeuil and Taneyhill, 2020; Li et al., 2020; Piacentino et al., 2020; Teddy and Kulesa, 2004). Our data combining *in vivo* and *ex vivo* approaches reveal an additional mechanism of communication whereby NC cells release EVs that can regulate behavior of placode cells. This is consistent with a recent study demonstrating that cranial NC cells produces EVs that are critical for their migration (Gustafson et al., 2022), albeit neither the contents nor the molecular function were addressed. Altogether, sEVs offer a delivery method for cell-to-cell communication in which miRNAs produced and released by donor cells are taken up by recipient cells where they can cause changes in gene expression (Thomou et al., 2017; Garcia-Martín et al, 2022; Valadi et al., 2007). Importantly, a recent study found that the miRNA population released in sEVs is distinct from the population found in the cells of origin (Garcia-Martin et al., 2022). Moreover, the authors demonstrated that miRNAs content enriched in sEVs differs by cell type and identified the presence of EXOmotifs and CELLmotifs, which may be recognized by RNA-binding proteins which participate in the specific sorting mechanism. In our study, we demonstrate that miR-203, presenting a consensus EXOmotif, is sorted into sEVs and can affect the expression of a sensor mRNA expressed in placode cells. This agrees with the fact that the loss of miR-203 function only affects trigeminal ganglia condensation when it is caused in placode cells. Thus, our study is one of the first to use an *in vivo* system to study the role of miRNA cargo in sEVs during intercellular communication in early development.

Finally, we demonstrate that neural crest-produced cytoneme-like structures contact placode cells. This structure participates in contact-mediated cell communication during early organogenesis, where cell protrusions are used to deliver signals (Mattes and Scholpp, 2018). Importantly, it was demonstrated that sEVs are transported along cytonemes (Gradilla et al., 2014). Based on this, we speculate that cytonemes produced by NC cells may enable a high local concentration of sEVs released near placode cells. Active engulfment of sEVs by placode cells ensures that a sufficient load of miRNAs reach the cytoplasm to mediate target inhibition. Altogether, the neural crest-placode interaction during sensory ganglion development offers an excellent *in vivo* model system in which to examine the mechanism by which selective miRNAs cargo is delivered into sEVs and transported to specific target cells.

## Acknowledgments

We thank all the authors and members in the Laboratory of Developmental Biology at the INTECH for their contribution and helpful discussions during the course of our study. We thank Dr. Alissa M. Weaver (Vanderbilt University School of Medicine, Nashville, TN, USA) for the pHluo construct. We are very grateful to the directors and students from “Escuela de Educación Secundaria Agraria de Chascomús” for providing fertilized eggs of excellent quality.

## Author contributions

Y.E.B. and P.H.S-M. designed, performed the experiments and wrote the manuscript with editing and input from all coauthors; E.S-V performed experiments; M.L.P. and H.U. contributed in the co-culture and time-lapse experiments; I.R and M.I.R contributed on the NTA analysis of sEVs; K.L.A.S and A.P-N. performed EM imaging and aided in their interpretation; M.E.B contributed in the discussion, experimental design and manuscript writing; N.d.M. contributed on the sEVs purifications, characterization, experimental design and manuscript writing.

